# Generalizable gesture recognition using magnetomyography

**DOI:** 10.1101/2024.09.30.615946

**Authors:** Richy Yun, Richard Csaky, Debadatta Dash, Isabel Gerrard, Gabriel Gonzalez, Evan Kittle, David Taylor, Rahil Soroushmojdehi, Dominic Labanowski, Nishita Deka

## Abstract

The progression of human-computer interfaces into immersive and touchless realities requires new ways of interacting with machines that are correspondingly intuitive and seamless. Among these are gesture-based systems that use natural hand movements to interact with and control digital devices. Today, these systems are most commonly implemented through the use of cameras or inertial sensors, which have drawbacks in environments that are poorly lit, in conditions where the hands are obscured, or for applications that require fine motor control. More recent studies have advocated for the use of surface electromyography (sEMG) to capture gesture information by sensing electrical activity generated by muscle contraction. While promising demonstrations have been shown, studies have also outlined limitations in sEMG when it comes to generalization across a population, largely due to physiological differences between individuals. Magnetomyography (MMG) is an alternative modality for measuring the same motor signals at the muscle, but is impervious to distortions caused by tissue, hair, and moisture; this indicates potential for lower variability caused by physiological differences and changes in skin conductivity, making MMG a promising generalizable solution for gesture control. To test this theory, we developed wristbands with magnetic sensors and implemented a signal processing pipeline for gesture classification. Using this system, we measured MMG across 30 participants performing a gesture task consisting of nine discrete gestures. We demonstrate average single-participant classification accuracy of 95.4%, rivaling state-of-the-art accuracy with sEMG. In addition, we achieved higher cross-session and cross-participant accuracy compared to sEMG studies. Given that these results were obtained with a non-ideal recording system, we anticipate significantly better results with better sensors. Together, these findings suggest that MMG can provide higher performance for control systems based on gesture recognition by overcoming limitations of existing techniques.

## 1 Introduction

Gesture recognition for human-computer interaction (HCI) aims to incorporate one of the most natural and intuitive forms of expression into how we control our everyday electronic devices. Gesture recognition has been widely studied and demonstrates promise in a variety of applications, including control systems, non-verbal communication methods, and physical rehabilitation (Guo et al., 2021; Mohamed et al., 2021; Yasen & Jusoh, 2019; Zaman Khan, 2012). Outside of the laboratory, commercially available virtual reality headsets, smart glasses, and smartwatches have already begun to employ control schemes using simple, discrete gestures. However, despite the appeal of gesture recognition, there has yet to be a ubiquitous solution due to difficulties in ensuring ease of use and generalizability.

Conventional approaches mostly fall within three categories: computer vision using cameras to track the hands, wired gloves to extract hand kinematics, or physiological measures like surface electromyography (sEMG) (Guo et al., 2021). Computer vision extracts joint positions using video streams from infrared and/or optical cameras. However, camera-based systems are limited by the need for an unobstructed view of the hands or fingers in good lighting and reliance on movement to detect activity (Abraham et al., 2015; Yasen & Jusoh, 2019; Zaman Khan, 2012). Wired gloves are composed of sensors that a person can don which tracks the position of each joint. Although highly accurate, wired gloves require the sensors to be placed directly on the hands reducing portability and dexterity, which precludes a number of potential use cases (Caeiro-Rodríguez et al., 2021; Dipietro et al., 2008).

In contrast, sEMG, a physiological measure of muscle activity, overcomes these limitations by directly detecting activity reflective of muscle contractions from the wrist or forearm using an arm-mounted device. With this approach, the hands can be positioned anywhere and free of sensors placed directly on the hands themselves. In addition, changes in isometric contraction, such as grip strength, can be properly measured allowing for an additional dimension of inputs (Merletti & Farina, 2016). Consequently, gesture recognition using sEMG is an active field of study and is employed for both research and commercial applications (Jaramillo-Yánez et al., 2020; H. S. Lee et al., 2022; Simao et al., 2019; Soroushmojdehi et al., 2022).

Despite these advantages, physiological variability between individuals as well as within-subject sensor displacements introduce major challenges in developing a generalized model of sEMG across a population (Eddy et al., 2024; Kaifosh & Reardon, 2024; Phinyomark & Scheme, 2018). Recent studies have shown notable progress with deep learning approaches: >95% accuracy between sessions of the same subject across 50 gestures (H. Lee et al., 2024), and >92% accuracy between 4800 subjects across nine gestures (Kaifosh & Reardon, 2024). Despite these encouraging results, the latter study concluded that their approach reaches a theoretical upper limit of around 95% accuracy across nine gestures using 16 channel sEMG on the wrist. This limit is likely due to differences in anatomy, physiology, and behavior between participants (Kaifosh & Reardon, 2024). Though groundbreaking to generalize across so many subjects, 5% error for a control system is likely to cause frustration and induce fatigue (Évain et al., 2016; Scheirer et al., 2002; Van De Laar et al., 2013). Thus, there is a clear need for a better muscle activity recording modality with less variability between participants to enable more generalized and accurate gesture recognition.

We propose magnetic sensing of muscles, or magnetomyography (MMG), as a novel approach for gesture recognition. MMG is a modality that has been demonstrated to provide a direct measurement of magnetic field generated by the electrical activity of muscles without requiring direct skin contact (Ding et al., 2022; Sometti et al., 2021; Yun et al., 2024). MMG is understudied due to the inaccessibility of recording systems, typically requiring extremely expensive magnetic sensors like superconducting quantum interference devices (SQUIDs) or optically pumped magnetometers (OPMs) (Arekhloo et al., 2023; Broser et al., 2018; Cohen & Givler, 1972). These sensors also need a magnetically shielded room to remove external magnetic interference for proper operation. However, recent advancement in sensor technology has improved accessibility of highly sensitive magnetic sensors and thus MMG, suggesting its potential applicability beyond research or medical settings (Labanowski et al., 2016; Yun et al., 2024; Zuo et al., 2020).

MMG has several advantages over sEMG for gesture recognition. Unlike sEMG, MMG does not rely on skin or tissue connectivity as the human body is largely permeable to magnetic fields (Cohen & Givler, 1972; Garcia & Baffa, 2015). Thus, MMG can measure muscle activity without distortions caused by the conductivity of tissue or electrode properties, both large contributors to the variability experienced by sEMG (Arekhloo et al., 2022; Klotz et al., 2023; Merletti & Muceli, 2019). In addition, many magnetic sensors for MMG can be miniaturized into chip-form factor allowing for high density of channels, further increasing the information content available for gesture recognition (Labanowski et al., 2016; Murzin et al., 2020; Zuo et al., 2020). As a result, MMG may provide less cross-participant and cross-session variability allowing for a more generalized model in gesture recognition.

To investigate this hypothesis, we collected multiple sessions of MMG recording from the wrist and forearm across 30 participants during a hand gesture task. First, we demonstrate that gestures can be accurately classified using MMG. Next, we assess cross-session and cross-participant classification to establish baseline variability. Finally, we present findings which show that the majority of the observed variability is due to the limitations of the available recording system rather than inherent to the modality of MMG. These results suggest that MMG is a promising modality for generalized gesture recognition.

## 2 Results

### 2.1 Gestures can be accurately classified with MMG

We collected multiple sessions of data across 30 participants with a total of 70 sessions with OPMs and sEMG electrode pairs (Figure 1a and Table 1). MMG signals were validated to actively reflect muscle activity by comparing the time domain signal and spectral density between MMG and sEMG (Figure 1b, 1c, respectively). We found similar SNR and spectral content between MMG and sEMG as shown previously (Yun et al., 2024).

**Table 1:** Participant Demographics and Session Details.

| Participant | Handedness | Sex | Age | Wrist Circ (cm) | Forearm Circ (cm) | # of Sessions | # of Bad Trials (per Session) | # of Bad Channels (per Session) |
| --- | --- | --- | --- | --- | --- | --- | --- | --- |
| 1 | R | F | 28 | 16 | 25.5 | 2 | 11,9 | 0,0 |
| 2 | R | M | 32 | 18.5 | 27.5 | 2 | 9,10 | 2,2 |
| 3 | R | M | 28 | 17.5 | 28 | 2 | 13,6 | 0,2 |
| 4 | Amb | M | 35 | 20.5 | 31 | 3 | 9,8,11 | 5,1,0,0 |
| 5 | R | F | 64 | 20.5 | 31 | 3 | 5,3,7 | 0,1,1,0 |
| 6 | R | M | 28 | 14.5 | 22 | 3 | 19,11,13 | 1,0,0 |
| 7 | R | M | 36 | 17 | 27 | 3 | 13,6,12 | 3,5,1 |
| 8 | R | M | 35 | 17 | 28 | 3 | 11,9,33 | 4,4,2,1 |
| 9 | R | M | 34 | 19 | 32 | 3 | 20,13,16 | 2,0,3 |
| 10 | R | M | 32 | 17 | 28 | 2 | 11,8 | 0,1 |
| 11 | R | M | 25 | 17 | 26.5 | 3 | 15,6,21 | 0,0,0 |
| 12 | R | F | 24 | 15.5 | 25 | 2 | 11,14 | 2,0 |
| 13 | R | F | 33 | 15 | 23 | 3 | 16,22,18 | 2,1,0 |
| 14 | R | M | 31 | 17.5 | 28.5 | 3 | 9,18,10 | 1,2,3 |
| 15 | R | M | 33 | 17 | 27 | 4 | 15,6,17,15 | 2,0,1,0 |
| 16 | R | M | 63 | 15 | 25 | 1 | 20 | 1 |
| 17 | R | M | 31 | 16.5 | 27 | 3 | 13,4,6 | 3,0,0 |
| 18 | R | M | 46 | 17 | 28.5 | 1 | 19 | 2 |
| 19 | R | F | 36 | 17 | 26 | 1 | 14 | 3 |
| 20 | R | F | 32 | 15 | 23 | 3 | 15,7,11 | 5,1,1 |
| 21 | R | F | 30 | 16 | 26 | 3 | 19,22,29 | 0,0,3 |
| 22 | R | M | 31 | 18.5 | 31 | 2 | 15,22 | 2,3 |
| 23 | L | F | 34 | 15 | 23 | 1 | 20 | 4 |
| 24 | R | M | 21 | 16 | 24 | 3 | 22,11,17 | 1,1,1 |
| 25 | R | F | 34 | 15 | 22.5 | 2 | 17,13 | 1,0 |
| 26 | R | F | 29 | 14 | 21 | 3 | 27,11,5 | 2,2,2 |
| 27 | R | M | 23 | 17 | 27 | 1 | 18 | 3 |
| 28 | R | M | 40 | 19 | 30 | 1 | 9 | 1 |
| 29 | R | M | 33 | 16.5 | 26.5 | 1 | 18 | 4 |
| 30 | R | M | 20 | 16 | 24.5 | 3 | 16,10,7 | 0,1,0 |
| Mean |  |  | 33.37 | 16.77 | 26.5 | 2.5 | 13.39 | 1.48 |
| Standard Deviation |  |  | 9.79 | 1.64 | 2.89 | 1.04 | 6.01 | 1.47 |

**Figure 1:**
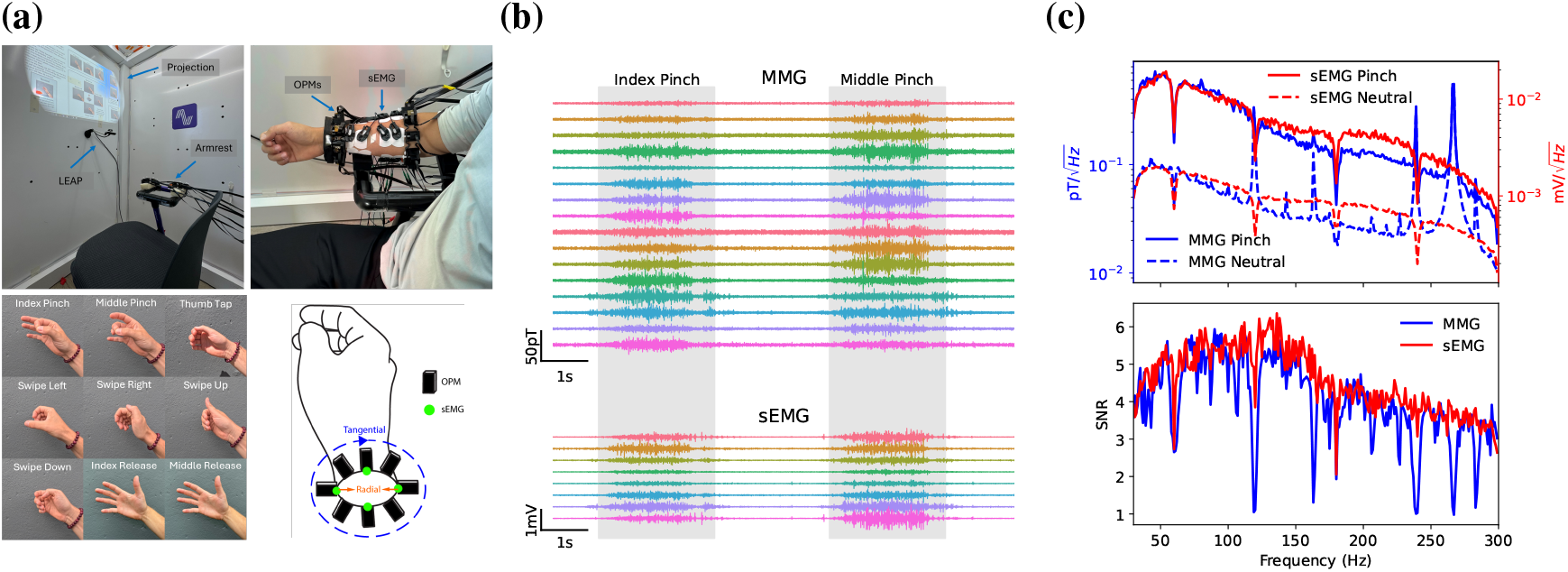
Experimental Setup. (a) Layout of magnetically shielded room (top left), the placement of OPMs and sEMG pairs (top right). Images of the 9 gestures presented to the participant to prompt a trial start (bottom left). Index and Middle Release were prompted immediately following the corresponding Pinch gesture. Placement of a band of OPMs and sEMG electrodes on the wrist (bottom right). Note the two directional axes of recording, tangential and radial. A similar band was placed on the forearm. (b) Example traces of 16 OPM channels (top 8 around the wrist, bottom 8 around the forearm) and all sEMG pairs during an Index Pinch and Middle Pinch trial. The signals were notched filtered for line noise (60 Hz harmonics) and bandpass filtered from 30 to 300 Hz. Note the similarity in timing and SNR between the two modalities. (c) Example average amplitude spectral density for a single OPM (blue) and EMG (red) channel on the dorsal side of the wrist (top). Solid lines show the average across all Index Pinch trials and dashed lines averages for all Neutral trials. SNRs of the two channels calculated by dividing the spectral density during Index Pinch by the spectral density during Neutral (bottom). Note the similarity between the spectral content of the signals.

We first established baseline single-session performance using MMG multivariate power frequency (MPF, see Methods) features with a Logistic Regression model including all 32 channels (16 at wrist and 16 at forearm) (Figure 2a). The median cross-validation accuracy across participants for 9 gestures was 95.4% [92.3% - 98.1% IQR]. All gesture types other than Index and Middle release gestures had >95% median accuracy. Index and Middle Release gestures performed significantly worse compared to all other gestures, with 87.2% [74.5% - 95.8% IQR] and 87.5% [77.5% - 95.7% IQR] accuracy respectively.

**Figure 2:**
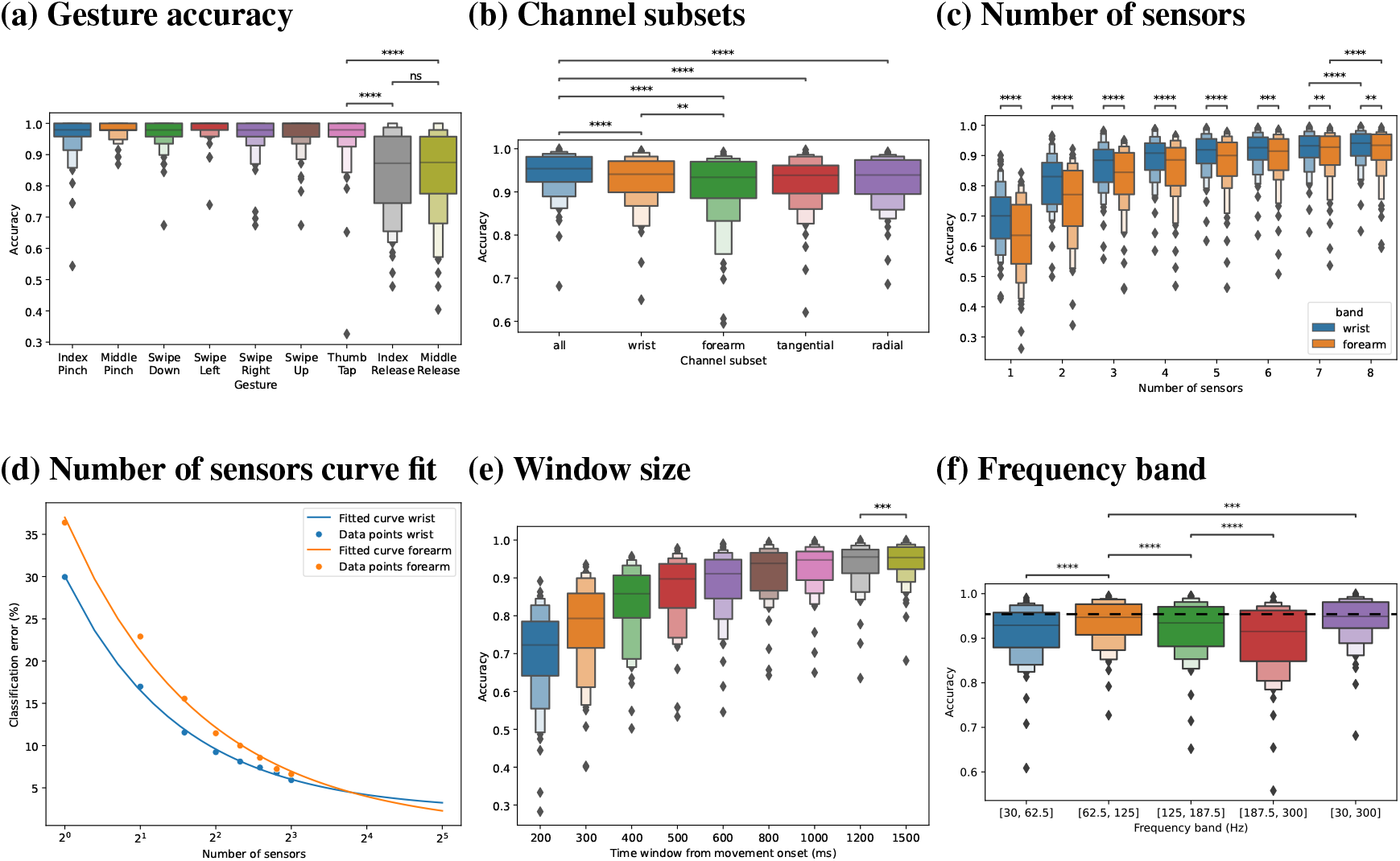
Single-session gesture classification. Distribution of single-session classification accuracies are shown for various analyses. Chance-level is 0.11. (a) Per-gesture cross-validation accuracy for single-session models using all channels. Release gestures had significantly worse performance than other gestures. (b) Cross-validation accuracy across various channel subsets. Best accuracies were obtained when using all 32 channels. Using channels at the wrist performed better than channels at the forearm. There was no difference between using just tangential or radial channels. (c) Cross-validation accuracy scaling with the number of sensors in each band. Performance improved steadily when increasing the number of sensors. Using the wrist sensors always had higher accuracy than the forearm sensors. For each session and sensor count we used multiple random samplings of sensors. (d) Curve fit to the median cross-validation classification error (100 - accuracy) from (c). The curve fit to the wrist channels was *Er* = 2.24 + 27.78*/N* ^0.96^ and to the forearm channels was *Er* = 0 + 37*/N* ^0.8^. (e) Cross-validation accuracy scaling with the window size used from movement onset. Performance improved steadily with larger window lengths with statistically significant differences between each subsequent window size. Distributions were over session-level cross-validated models trained on different window lengths from movement onset. (f) Cross-validation accuracy across various frequency bands. The horizontal dashed line denotes accuracy obtained with the entire bandwidth. The 62.5-125 Hz band contained the most information content with every other frequency band having less information. Distributions were over session-level cross-validated models trained on the respective frequency band.

Next, we compared classification accuracies between subsets of channels to better understand the significance in channel position between the wrist and the forearm as well as axis of sensitivity in classification accuracy (Figure 2b). Using all channels, the median cross-validation accuracy was 95.4% [92.3% - 98.1% IQR], while wrist-only channels was slightly lower at 94.1% [89.9% - 97.1% IQR], and forearm-only channels was 93.4% [88.5% - 97.0% IQR]. All comparisons between these 3 channel subsets were statistically significant (p<1e-3, Bonferroni corrected for 5 comparisons) showing that while signals from the wrist potentially have more distinguishable features specific to gestures compared to the forearm, the forearm may provide additional information not present in the wrist. Using just tangential or radial channels achieved similar performance, with 93.9% [89.6% - 96.1% IQR] and 93.9% [89.5% - 97.4% IQR] accuracy respectively. The differences between the axes were not statistically significant. We present confusion matrices for each training in Supplementary Figures S1-S4.

Additionally, we sought to investigate how the number of sensors affects classification performance. To that end, we trained the same Logistic Regression model with different number of sensor inputs from the wrist and forearm bands separately. In each case, we sampled 16 random sensor subsets, except in the case of 1 and 7 sensors where only 8 total different variations are possible. For each sensor we used both axes, as we previously determined that there is no large difference between the two. Figure 2c shows classification performance with respect to the number of sensors used to train the model. Across any number of sensors the forearm sensors resulted in statistically lower performance compared to wrist sensors (p<1e-3, Bonferroni corrected for 8 comparisons). Though slowly reaching an asymptote, performance continued to increase significantly even when going from 7 to 8 sensors for both wrist and forearm bands (93.5% to 94.1% accuracy and 92.7% to 93.4% accuracy respectively, p<1e-10).

With the assumption that the relationship between the classification error and number of sensors follows a power law, we fit a curve in the form of 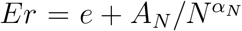 to the classification error (*Er*) with respect to the number of sensors (*N*)(Figure 2d). The curve suggests that performance should continue improving drastically with increase sensor count. Interestingly, the performance of the forearm band overtakes the performance of the wrist band at around 14 sensors, though this may potentially be due to the oversimplification of the curve fit.

Next, we explored the effect of MPF feature window length on classification performance. The model was trained on varying window lengths following movement onset, with MPF features calculated using two non-overlapping windows (Figure 2e). All consecutive pairs of window lengths, even when going from 1200 to 1500 ms, were significantly different (p<1e-2 for 1200 ms vs 1500 ms, p<1e-5 for all other pairs, Bonferroni corrected for 8 comparisons). Only a 500 ms window was required to achieve close to 90% median accuracy (89.7% [82.1% - 93.7% IQR]). Interestingly, Index and Middle Release gestures were typically better classified using a smaller window compared to the full trial length whereas Index and Middle Pinch gesture classifications were the most affected by a reduced window length (Supplementary Figure S2). However, we also found large differences between participants in how window size affects classification accuracy for specific gestures suggesting additional data is necessary to draw conclusions on the impact of window length across different gestures.

Finally, we investigated which frequency bands contained most of the gesture-related signals. To this end, we trained the model on each frequency bin used in the MPF computation: 30-62.5, 62.5-125, 125-187.5, and 187.5-300 Hz. Figure 2f shows that the highest accuracy was achieved using the (62.5, 125) Hz band (94.7% [90.8% - 97.6% IQR]), almost as high as when using the entire bandwidth (95.4%). Accuracy was significantly lower in all other frequency bands (p<1e-5, Bonferroni corrected for 4 comparisons); however, we were still able to achieve high accuracy even when using the 187.7-300 Hz band (91.5% [84.8% - 96.2% IQR]) which was unexpected due to the limited bandwidth of the OPM sensors.

### 2.2 MMG is generalizable across sessions and subjects

#### Cross-session classification

To first determine how well MMG can perform from session to session on the same participant we performed cross-session analyses, in which the model was trained on one or two of three sessions. When using one training session a second session was used for validation and the third session for test. When using two training sessions, validation was sampled randomly from the 2 training sessions (10% of trials) and the third session was used for test (3 folds in total). On each fold we trained a CNN+LSTM model (see Methods) on time-aligned data (1.5s trial length) on all channels as well as wrist-only and forearm-only channels. We used data from 17 participants that had at least 3 sessions and we report mean test session accuracy over each train-test choice for each participant.

Cross-session performances is shown in Figure 3a. Using 2 training sessions instead of 1 increased performance substantially, achieving 71.5% [61.8% - 78.6% IQR] compared to 58.6% [47.9% - 72.5% IQR] (p<1e-7). Cross-session generalizability was additionally tested using just the wrist or forearm channels which also showed significant increase in performance when using 2 training session compared to just 1 training session (p<1e-7). Generalization performance between the wrist and forearm channels was not significantly different from one another when using 1 or 2 training sessions. Unsurprisingly, Using all channels performed better than either subsets of channels (p<1e-3, Bonferroni corrected for 4 comparisons). Similar to single session classification, Index and Middle Release gestures were the most misclassified (Supplementary Figure S3).

**Figure 3:**
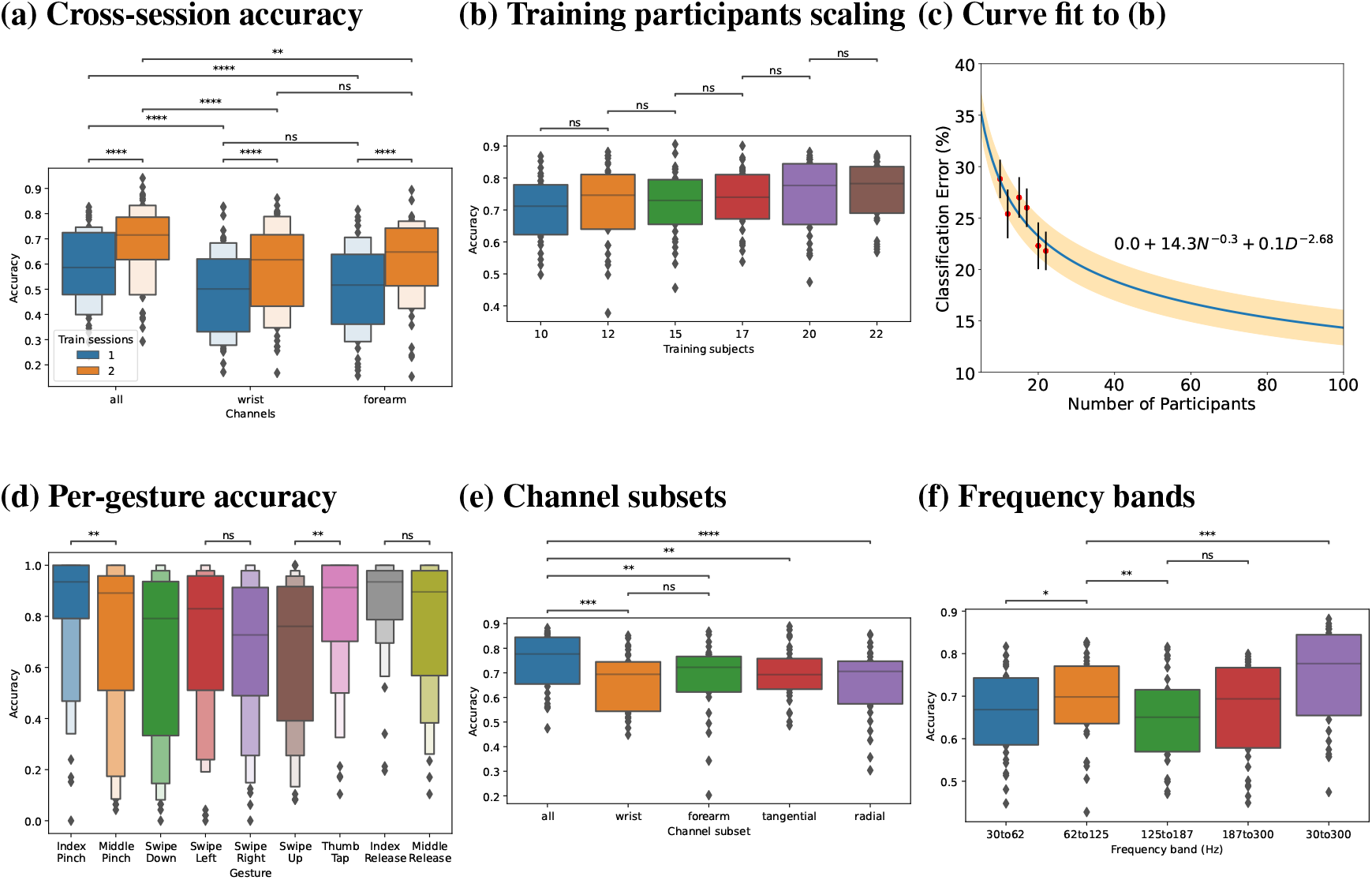
Cross-session and cross-participant results. Chance-level is 0.11. (a) Cross-session accuracy across test sessions. Using 2 training sessions instead of 1 substantially improves performance, as well using all channels compared to a subset of channels. Distributions consist of sessions across all participants. There was no difference when using wrist-only or forearm-only channels. (b) Cross-participant accuracy across test participants depending on the size of the training set. Performance improves steadily when adding more participants to the training set. Distributions are over the mean (across sessions) accuracies of each test participant. (c) Curve fit to the median classification errors (100-accuracy) from (b) and extrapolated to 100 participants. The following parameters for the best curve fit was found *e* = 0.0, *A*_*N*_ = 14.33, *α*_*N*_ = 0.3 (Equation 1) Black bars denote SEM ± around the median, and the orange band denotes the area between the curve fit to the upper and lower SEM values giving estimates of the error around the median. (d) Per-gesture accuracy for the cross-participant model using all channels. Swipe Right and Swipe Up have the worst performance. (e) Cross-participant accuracy across test participants using various channel subsets. Using all channels has substantially higher performance than all other combinations. Distributions are over the mean (across sessions) accuracies of each test participant. (f) Cross-participant accuracy across test participants using various frequency bands. Best generalization performance is achieved with the 62-125Hz and 187-300Hz ranges, but this is lower than using the whole 30-300Hz range.

#### Cross-participant generalization

To determine generalizability of a model across participants we performed cross-participant classification using all available sessions for each participant. In order to provide robust results in each of the following analyses we trained the model 30 independent times, 1 for each test participant, where the remaining 29 participants were randomly split into train and validation sets. We trained the CNN+LSTM model (see Methods) on non-time-aligned data using the full trial length (2.5s) with all 32 channels, and used a leave-1-participant-out testing setup by sampling the train and validation participants randomly for each test participant.

Figure 3b shows the effect of the number of participants used in the training data on the generalized model performance. Though we did not find a statistically significant difference between consecutive pairs of numbers of training participants, this was likely due to the limited number of repetitions we performed for each count. Nevertheless we did find a general trend with increasing accuracy with increasing number of training participants, reaching a peak of 78% [69.0%-83.5%] accuracy with 22 participants.

To better quantify the relationship, we fit an exponential curve of the following form to the median accuracies of the training subsets following (Kaifosh & Reardon, 2024):

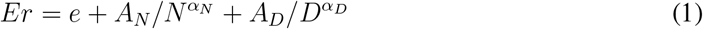

Where *Er* is the classification error measured in percentage, *N* is the number of participants measured in hundreds, *D* is the model size (millions of parameters). *e, A*_*N*_, and *α*_*N*_ are variables to fit a power law defining irreducible error (i.e. expected error with infinite number of participants) and the rate of change with respect to the number of participants. *A*_*D*_ and *α*_*D*_ are parameters corresponding to model size – as we trained on a single model size, we used values found in (Kaifosh & Reardon, 2024): *A*_*D*_ = 0.1, *alpha*_*D*_ = 2.68. Calculating the fit with our results we found the following parameters: *e* = 0.0, *A*_*N*_ = 14.33, *α*_*N*_ = 0.3 (Figure 3c). Except *e*, which is the irreducible error, these parameters are close to those reported in (Kaifosh & Reardon, 2024). Though we see extremely low irreducible error, this may be an artifact of extrapolation with not enough participants and warrants further investigation.

Finally, to assess whether certain gestures were easier to generalize we show the per-gesture differences in performance for the 20-training-participant analysis in Figure 3d. Swipe Right and Swipe Up gestures had the lowest performance, while Index Pinch and Index Release had the best performance. There were significant differences between Index and Middle Pinches, but not between Releases. Swipe Left was significantly better than Swipe Right. All tests were Bonferroni corrected for 4 comparisons. Interestingly, the Index and Middle Release gestures that had the lowest performance for single subject performance were classified well in generalized models, even compared to other gestures.

#### Channel subset analysis

Similar to single subject classification, we trained the CNN+LSTM model on all channels, as well as just wrist, forearm, tangential, and radial channels for cross-participant analysis (Figure 3e). Across each of these trainings we matched the train-validation splitting for each test participant by using the same random seed. Using all channels performed better than any subset of channels (77.7% [65.5% - 84.4% IQR], p<1e-5, Bonferroni corrected for 5 comparisons). Forearm channels (72.3% [62.2% - 76.6% IQR]) performed slightly higher than wrist channels (69.4% [54.4% - 74.5% IQR]), but not statistically so. Tangential and radial channels had similar performances (69.3% [63.3% - 75.8% IQR] and 70.6% [57.4% - 74.7% IQR] respectively), and were not significantly different from on another.

Out of the 30 total participants we found that 3 participants had higher mean accuracy with wrist channels, and 3 participants had higher mean accuracy with forearm channels. The confusion matrices summed across all test participants, of 14 sessions with highest and lowest test accuracy, as well as of wrist or forearm channels are shown in Supplementary Figure S4. There were no consistent differences between which gestures were most confused. Overall, this demonstrates the large variability between participants when fitting a generalized model.

#### Frequency bands

Finally, as we did for single-participant trainings, we investigated which frequency bands contributes the most to generalizability across participants. We performed similar analysis as before using all channels on data that was bandpass filtered using the following ranges: 30-62, 62-125, 125-187 187-300 Hz (Figure 3f). Trends are similar to the single-participant case, except that the 187-300 Hz frequency band performed almost as well as the 62-125 Hz band. This suggests that high frequency content, even if containing less overall information, may be more generalizable across participants. Analyzing each pairwise comparison, we found that the only significant difference was between the 62-125 Hz and 125-187 Hz bands (p<2e-2). All individual frequency band trainings were <70% median accuracy and significantly lower than using the entire 30-300 Hz bandwidth (p<1e-3, Bonferroni corrected for 10 comparisons).

### 2.3 Sources of variability

#### Accuracy over the duration of each session

Possible causes of lower performance in certain sessions or participants include the shifting/rotation of the OPM band(s), as well as changes in participant position in the shielded room. OPM sensors operate by detecting and counteracting a bias field with built-in coils which can only be calibrated at the beginning of each session, but the movement changes the bias field the OPMs are subjected to, thus reducing the efficacy of the coils. An increase in the bias field can present itself as increases in movement artifacts, as there are larger fields the sensor is now moving through, as well as saturation of the sensor in worst case scenarios. In addition, increased bias fields reduce linearity of the sensors. If movement from initial baseline increases as the recording goes on, we would expect performance to decrease over time.

We selected the best 14 and worst 14 sessions according to accuracy (for each channel-subset cross-subject training), and ran a sliding window of 20% of the number of trials over the per-trial accuracies. In Figure 4 we plot accuracy over consecutive trials within the recording. While the trends are minimal, for example in the all-channels case (Figure 4a), the low-accuracy sessions do show an overall decrease in performance with respect to recording duration. Interestingly, the high accuracy sessions in the forearm-only training also show a similar trend, suggesting this only accounts for a part of the differences between low- and high-performing sessions.

**Figure 4:**
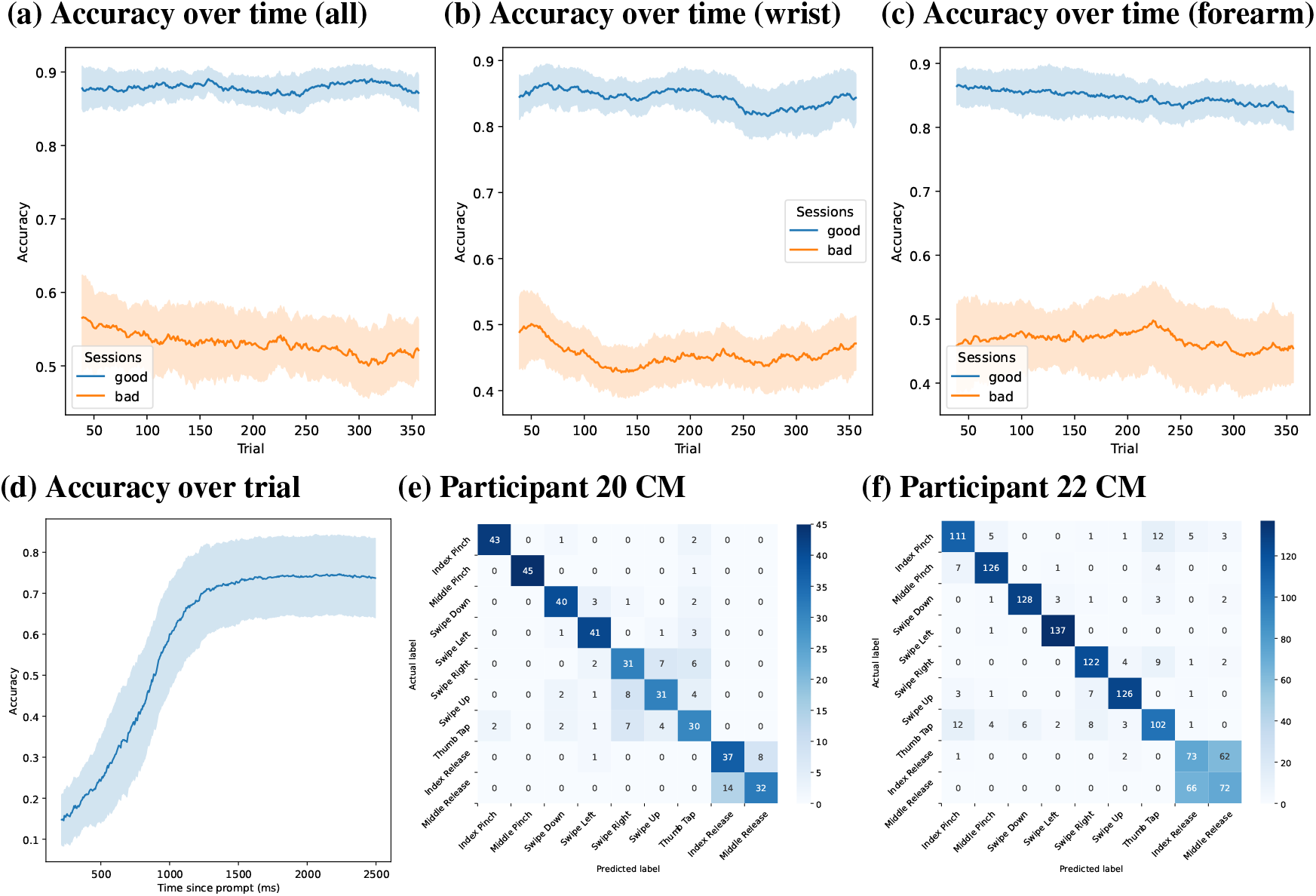
Variability across recording duration, trial timing, and between participants. (a-c) Cross-subject accuracy over the recording duration for 3 channel subsets; all, wrist, and forearm. All cases show a decrease in accuracy over the recording duration except the all-channels *good* sessions. *good* refers to the top 14, and *bad* refers to the bottom 14 sessions according to test accuracy. The horizontal axis denotes the ordered trial index within the recording. Note that a sliding window of 20% of all trials (about 80 trials, depending on the session) was used. (d) Cross-subject accuracy over the entire trial length showing median accuracy and standard deviation across participants. Accuracy increases rapidly in the beginning of the trial plateauing around 1400ms. (e-f) Aggregated confusion matrices of 2 participants with low performance. These were obtained from MPF+logreg models trained on individual sessions (using all channels). The confusion matrices show that there are differences between participants in terms of which gestures are poorly classified.

To quantify these measures, we calculated a linear fit for each session partition (i.e. mean accuracy across top-14 and bottom-14 sessions) and channel subset (Bonferroni corrected for 6 comparisons). While the linear trend slope is very small, we still found significant (p<1e-6) downward trends in almost all cases (not only the ones highlighted above). Thus, both high-accuracy and low-accuracy sessions show this trend. The only exception was the high-accuracy sessions for the all-channels training which showed a significant (p<1e-2) positive trend. We also correlated each session’s accuracy slope with the overall accuracy value of that session, to see if low-accuracy sessions show a larger downward trend, but we did not find a significant relationship (p>0.1).

#### Window size

As with single-session classification, we wanted to assess performance with respect to the window size for cross-subject trainings. To this end we trained a modified CNN+LSTM on non-time-aligned data (2.5 second trials) with an extra data augmentation. This involved randomly cropping the start of each trial up to a maximum of 1000 ms. Testing trials were always 2.5 seconds long. To enable a single CNN+LSTM model to be able to predict across different window lengths, we included the output of each LSTM timestep starting from roughly 200ms in the loss function. This means that output predictions based on the first 200ms were not optimized in the loss. We trained this model on the same splits/folds as our original 20-training-participant all-channel model.

After training, we generated predictions and computed accuracy at each timestep, resulting in Figure 4d. Since the CNN+LSTM was trained on the entire trial (starting from the prompt) early timesteps have low accuracy, as there is little information present. Performance increases rapidly, plateauing around 1400ms. We calculated repeated measures ANOVA across timepoints to assess statistical differences with respect to time and found p<1e-4. Then we ran pairwise comparisons (Wilcoxon signed rank tests, Bonferroni corrected for the number of timepoints) between each timepoint and the final timepoint. We found that the first time the p-value goes above 0.05 is at 1410ms, and thus this is the point where accuracy stops increasing significantly. This shows that most information necessary to classify gestures across participants is present in the first 1400ms following the prompt. Taking into account the time to movement onset the actual information content window is likely much smaller.

#### Confusion matrices per subject

There is considerable variability in which gestures have better or worse performance between participants. In Figure 4e and 4f we show individual confusion matrices from the single-session MPF+logreg trainings. We selected the two participants with lowest accuracy to illustrate variability. While participant 22 has almost chance-level performance in the release gestures, participant 20 has much lower performance in the swipe gestures compared to others. We investigate the confusion matrices of these two participants for wrist-only and forearm-only trainings in the Supplementary Material.

## 3 Discussion

We demonstrated MMG measured at the wrist and forearm as a modality for hand gesture recognition. We first showed that MMG can be used to accurately classify between 9 distinct gestures. Our results further revealed that MMG generalized across sessions as well as across participants. The high frequency components were significant for both single- and cross-participant classification, suggesting the lack of distortion from the body to magnetic fields may be a contributing factor. Finally, we expanded upon the factors that contribute to variability observed in the dataset, a large part of which is due the limitations of the sensors used in the study.

### 3.1 MMG can be used to accurately classify hand gestures

We demonstrated high classification accuracy across 9 gestures with a median of >95%, rivaling state-of-the-art classification performed with sEMG (Côté-Allard et al., 2019; Kaifosh & Reardon, 2024; Xiong et al., 2021), despite a high rate of saturated trials and sensors (>13 trials and 1.48 saturated sensors per session on average, Table 1). The only previous attempt at gesture recognition using MMG relied on 8 channels along the forearm to classify between 3 states: index finger flexion, little finger flexion, and neutral (Greco et al., 2023). The authors were able to successfully distinguish between the gestures, but were not able to reach >90% accuracy. In addition, they showed that sEMG provided more accurate results despite only having 4 channels. Even when restricting the number of channels to 8 channels we saw similar or higher accuracies while having a larger gesture set (Figure 2c).

One large difference from Greco et al., 2023 was our much larger bandwidth, ranging from 30-300 Hz compared to 25-100 Hz. By performing classification using separate frequency bands (Figure 2f), we showed that high frequencies, despite being limited by OPM frequency response, provided significant information for classification. This also suggests that MMG’s ability to capture more high frequency power than sEMG due to the lack of tissue attenuation (Yun et al., 2024) will become increasingly important as new sensor technologies improve bandwidth.

In addition, we observed differences in the MMG signals obtained from the wrist compared to those from the forearm. Though intuition says the forearm should provide better signals due to higher muscle mass, sEMG signals have consistently shown higher classification accuracy when recorded at the wrist compared to the forearm, especially with deep learning approaches (Botros et al., 2022; He et al., 2024). This has been attributed to both the higher density of muscle groups at the wrist, allowing for higher information per channel, as well as the presence of muscles for fine finger motion that are not present in the forearm (Botros et al., 2022; He et al., 2024; Olsen et al., 2023). We saw similar results with MMG, showing significantly higher accuracy when recorded at the wrist compared to the forearm, demonstrating MMG is also viable for a wrist wearable device.

Furthermore, we were able to assess the impact of the axis of sensitivity of MMG on gesture classification. As a magnetic field curls around a current (Biot-Savart law) different vector components are measured by the different axes of sensors. Previous studies have shown that radial and tangential components have different SNRs depending on the muscles measured, with wrist and forearm measurements typically providing larger signals in the radial direction (Broser, Marquetand, et al., 2021; Broser, Middelmann, et al., 2021; Sometti et al., 2021; Yun et al., 2024). However, we saw no significant difference in gesture classification accuracy between radial and tangential components. Besides these two components, previous studies suggest the axial axis – orthogonal to both the tangential and radial directions – may have additional signal components due to the non-homogeneity of muscle fiber orientations (Broser, Marquetand, et al., 2021). Additional investigation is necessary to determine the potential benefits of obtaining multiple components of MMG from the same location.

### 3.2 MMG can be generalized across sessions and participants

We were able to generalize gesture classification using MMG both across sessions and across participants at higher accuracy than previously reported with sEMG (Côté-Allard et al., 2019; Du et al., 2017; Kaifosh & Reardon, 2024). This is likely due to various factors impacting sEMG session variability – such as humidity, skin dryness, skin/fat thickness, and exact placement of the sensors – being irrelevant to MMG as the human body is largely transparent to magnetic fields (Arekhloo et al., 2022; Nath et al., 2022). We were able to achieve our results despite using a non-ideal recording system with limited bandwidth and more than one saturated channel per session on average. As such, intrinsic MMG variability is likely to be even lower. In addition, when fit to the cross-participant accuracy we observed the non-generalizable cross-participant error was 0.0% (Figure 3c). Though additional investigation is necessary to confirm these findings, especially as we extrapolate to higher numbers of participants, the results suggest MMG may be an excellent modality for enabling out-of-the-box usage of a gesture recognition device.

A major source of variability we were able to discern was decrease in classification accuracy over time throughout the course of a session. As the train and test trials are randomly chosen, this suggests the latter trials typically have poorer signals. These changes are not likely due to simple rotation of the wristband, which would cause inconsistencies over time but not a monotonic change in classification accuracy. Instead, the variability is likely due to the motion of the participant in the shielded room causing movement of the sensors, introducing different noise profiles and saturating channels. Additional evidence, such as wrist sensors which are more likely to have moved performing worse than forearm sensors for generalization, also suggest that sensor saturation is a large contributor to classification performance.

Though denoising techniques exist, including use of accelerometers or camera based systems to track the sensors (Holmes et al., 2018; Seymour et al., 2021, 2022), gradiometers or synthetic gradiometry using additional sensors (Fife et al., 1999; Limes et al., 2020; Seymour et al., 2022), and various source separation methods (de Cheveigné, 2010; Taulu & Simola, 2006; Uusitalo & Ilmoniemi, 1997), they rely on offline processing or exact positioning of the sensors. To best emulate a wrist wearable we opted to not apply these techniques and instead rely purely on directly recorded signals with simple time domain preprocessing. Thus, either a recording system that has larger dynamic range and allows for participant movement or a real-time denoising approach are necessary to enable an MMG wearable.

In addition, we sought to determine if any metric was correlated to generalization performance. We found that though the number of bad trials (trials with multiple channels saturated) and bad channels (channels saturated throughout the duration of the session) were correlated with single-session classification accuracy, it was not correlated with cross-participant accuracy (Supplementary Figure S6b). Single-session classification accuracy itself was also not correlated to cross-participant accuracy (Supplementary Figure S6c). A large contributor may be physiological or task performance differences between participants (Supplementary Figure S8), but we also found individual sessions of the same participants could perform quite differently, suggesting that inter-session differences, including sensor positioning, how each trial is performed, and perhaps the noise profile difference between days, may also impact our cross-participant generalization. Transfer learning with subject calibration or subject embedding to enable transfer learning (Côté-Allard et al., 2019; Csaky et al., 2023; Kaifosh & Reardon, 2024; Soroushmojdehi et al., 2022), or better handling of poor data to catch incorrectly performed trials and high noise channels may enable better generalization with fewer participants. In addition, further analysis into latent space projection of the datasets (Supplementary Figure S5), potentially by exploring representations of various layers within the network (Kaifosh & Reardon, 2024), will help elucidate the exact differences between sessions and participants.

Finally, band passing the signals to different frequency bands showed 187-300Hz performed as well as 62.5-125Hz, suggesting that high frequency content is a significant contributor to not only classification accuracy but also for generalization. MMG’s ability to capture more high frequency power than sEMG (Yun et al., 2024) is likely a large factor in the observed better generalization performance.

### 3.3 Limitations and future directions

The OPMs used in this study, though providing incredibly sensitive magnetic recordings of muscle, are prone to saturation even in a magnetically shielded room (MSR). OPMs use biasing coils to remove DC fields to improve dynamic range (Osborne et al., 2018). However, movement of the sensor body such that the bias field changes can result in failure of the coils and saturated channels. The coils are also prone to introducing crosstalk between channels and sensors which prevented placement of a higher density of sensors just on the wrist. Additionally, the OPMs have limited bandwidth, with decreasing sensitivity beginning at 200 Hz and a 300 Hz hardware filter for the specific system used in the study. As a result, OPMs are not suitable for non-laboratory applications.

A small, sensitive magnetometer that allows for high proximity to the body as well as each other with larger bandwidth and dynamic range would enable much more stable recordings. Production of sensors that fulfill these requirements is an active field of study, with many promising results in applications even outside of MSRs (Labanowski et al., 2017; Limes et al., 2020; Zuo et al., 2020). A study utilizing these sensors would truly demonstrate the capabilities of MMG by decoupling the modality with the limitations of the recording system.

As this study is one of the early forays into applying MMG for gesture recognition, additional investigation is necessary to verify both the findings and additional characteristics of MMG. Reproducing previous sEMG work such as more complex or larger sets of gestures (Kaifosh & Reardon, 2024; Linderman et al., 2009; Wu et al., 2016) or continuous joint tracking (Ameri et al., 2019; Liu et al., 2021; Xia et al., 2017) will further elucidate the similarities and differences between sEMG and MMG. In addition, further research specific to MMG - exploring combinations of sensing axes, assessing effects of differences in physiology introducing changes in distance between muscle fibers and the sensor, how muscle size and density affects MMG signal amplitude, as well as deeper examination of the high frequency components that seem to be critical for gesture recognition - will provide a better understanding of how to apply MMG.

## 4 Acknowledgements

The authors would like to thank Dr. Sergey Stavisky for his insightful feedback and the participants of the study for volunteering their time.

## 5 Methods

### 5.1 Experimental design

#### 5.1.1 Magnetically shielded room

All experiments were performed inside a magnetically shielded room (MSR; Magnetic Shield Corporation, MuROOM) with inside dimensions of 1.3×1.3×2 meters and contamination from ambient noise sources. The shielded room had around 25,000 fold attenuation of residual fields at DC and up to 8000 fold attenuation at AC. The room was degaussed (demagnetized) by applying a decreasing sinusoidal field using embedded coils prior to each recording to remove magnetization of the walls and guarantee maximum rejection of ambient fields. A projector mounted on the outer MSR wall displayed a screen on the inside of the MSR through a hole in the wall.

#### 5.1.2 Sensors

We used the QZFM Gen-2 (QuSpin, Inc.) optically pumped magnetometer (OPM) sensors for MMG measurements. The OPM sensors were housed in a 3-D printed band containing 8 sensors each. Two sensor bands, one for the wrist and one for the forearm, were affixed to the participant’s arm. Each OPM was sensitive along two orthogonal axes, and were positioned such that one axis was tangential to the arm, and the other radially into the arm (Figure 1a). The OPMs recorded from both axes of each sensor for each recording. Throughout the manuscript we refer to data from individual axes as channels, as opposed to sensors. Thus, data collection involved 16 OPM sensors (8 per band) with 2 axes per sensor, totalling 32 MMG channels. Multiple size circumference bands were printed to accommodate variations in the circumference of the participants wrists and forearms. The wrist sensor band was placed 4cm up the arm from the wrist joint, and the forearm sensor band was placed at one third the length of the forearm down from the elbow. The sensors sat in an circular orientation, where one sensor was seated at the mid-line of the anterior face of the forearm and the other 7 sensors sat equidistant from each other around the circumference of the arm (Figure 1a).

We simultaneously recorded surface electromyography (sEMG) from each subject with single-use, bipolar, MRI safe, pre-gelled sEMG electrodes with non-ferrous contacts (EL508, Biopac Systems Inc., 10 mm gelled area diameter) and with non-ferrous leads (LEAD108, Biopack Systems Inc.). 16 sEMG electrodes were also adhered to the participant’s arm. The 16 electrodes were paired into 8 bipolar channels with 2.5cm center-to-center inter-electrode distance, 4 pairs per wrist and forearm. For both the wrist and forearm, two electrode pairs were placed at the mid-line of the anterior and posterior faces of the forearm, and two additional pairs were placed equidistant between those initial pairs, covering the forearm between the two OPM bands at approximately 90 degrees apart from each other. sEMG was used as a control to validate the MMG data.

Finally, we used an infrared-based hand tracking camera (Leap Motion Controller 2, Ultraleap) which allowed investigators to monitor the participant’s gestures from outside the MSR and simultaneously collect supplemental spatial hand position data during the gesture task.

#### 5.1.3 Data acquisition

The OPM and sEMG sensors provided analog signals which were digitized and streamed using 16- or 24-bit NI-DAQ ADC modules (one NI-9202 and two NI-9205 modules in a cDAQ-9185 chassis, National Instruments) at a sampling rate of 2 kHz per channel. The data from the Leap Motion Controller was streamed through Unity (a game engine) and directly saved in a separate file at around 100 Hz sampling rate.

#### 5.1.4 Gesture task

The task consisted of 12 distinct hand gestures: Clench, Extend, Flex, Index Pinch, Index Release, Middle Pinch, Middle Release, Swipe Right, Swipe Left, Swipe Up, Swipe Down, and Thumb Tap. Three of the gestures, Clench, Extend, and Flex, were used to ensure signal fidelity and not included in any classification analyses.

All swipe motions were performed with the thumb. The participant was cued to perform a specific gesture and then cued again to return to a “Neutral” position with the exception of Index and Middle Pinch which were immediately followed by Index or Middle Release. Each cue consisted of the image and name of the gesture as well as the number of trial repetition. For the Index and Middle Pinch gestures, the participant was cued to extend all of their fingers before returning to a neutral position to simulate the act of completing the pinch. The order of gestures was pseudo-randomly chosen and each gesture was performed for 50 trials.

A single session consisted of 50 trials of each gesture. Individual trials lasted 2.5 seconds and participants were instructed to hold their final position until the neutral cue was presented. Neutral positions also lasted 2.5 seconds between each trial. Each session began with 30 seconds of rest in which the participant held the Neutral position to obtain a baseline recording.

### 5.2 Participants

#### Recruitment

The data in this study was collected from 30 participants, ranging from 21 to 64 years old (Table 1). There were 10 female and 20 male participants; 28 participants were right handed, 1 was left handed, and 1 was ambidextrous. Our recruitment and data collection for human research followed IRB approved protocols (Advarra). Participants were recruited via an online form sent to their email, and interested individuals were scheduled and consented before their first data collection session. Personally identifiable information (PII), including demographic information (name, email address, phone number) and biometric information (bio-signals, arm length, age, weight, height, and additional information per the IRB) were housed and recorded separately following IRB approved confidentiality protocols. PII access was restricted to the PI only, and investigators were blind to PII data. Participants were instructed to undergo three data collection sessions, in which they performed the task over the period of roughly 40 minutes per session. Participants were compensated after each session with a gift card.

#### Participant briefing

All participants that consented to research were given a handedness questionnaire (Oldfield, 1971) to determine their dominant arm, as well as a metal screen to ensure that no metallic objects in or on the participants’ body entered the MSR. Participants were given detailed instructions on how to perform the task.

To get familiarized with the task, participants practiced the task for a few trials prior to the start of the experiment. Using a printout of the same gesture images that were displayed as cues during the task, the overseeing researcher demonstrated each gesture with the respective image, then performed the gestures alongside the participant. During this overview, the researcher guided the participants’ gestures, helping to ensure the individual’s consistency. These tips include real-life analogies and equivalents for each gesture all while the researcher simultaneously demonstrated the expected gesture with each instruction.

Finally, participants were asked to remove their shoes and change into scrubs and swept with a metal detector to fully confirm the absence of ferrous materials before they began each session. They were then seated in a non-ferrous chair inside the MSR. All sensors were subsequently affixed to their right arm which was then placed on a plastic armrest with foam padding for the duration of each session.

### 5.3 Data analysis

#### 5.3.1 Preprocessing

A common preprocessing pipeline was applied to the MMG data of all sessions for classification of gestures. The number and type of preprocessing steps may differ between different analyses, and these are specified in the respective sections. The pipeline consisted of the following steps.

##### Filtering

Raw data was bandpass filtered between 30-300 Hz, with a 5th order Butterworth filter. Then, a custom automatic peak finding algorithm was used to find noise peaks in the power spectral density (PSD) of each session which were subsequently removed with a 5th order Butterworth bandstop filter. The algorithm consisted of a sliding window along the PSD of each channel, where if within a certain frequency bin the z-scored power exceeded a threshold of 5, it was identified as a peak. The statistics for z-scoring were calculated based on a surrounding window of 20Hz. The algorithm was run in a 2-pass mode, removing peaks detected in the first pass and then running a second pass, again removing newly detected peaks. Finally, all data was resampled to 1000Hz.

##### Bad channel detection

After filtering, bad channels were automatically detected using the generalised extreme studentized deviate (ESD) test^1^ (Rosner, 1983), with a significance level of 0.5, and the maximum number of channels to mark as bad was set to 25% of all channels. The algorithm was applied separately to OPM and EMG channels. For single-session classification these channels were removed, but not for cross-session and cross-subject classification. We kept all channels for the latter two analyses to be able to match channel order between sessions.

##### Normalization

Channels were independently normalized to 0 mean and unit variance.

##### Epoching

Data were separated into 2.5 second-long epochs based on gesture prompt onset and offset. The generalised ESD test was used to automatically remove outlier epochs. This was calculated separately for each gesture type. The algorithm was run separately for OPM and EMG channels, and epochs identified as bad trials across either channel type were removed. The significance threshold was set to 0.1, and the maximum number of epochs to remove was set to 5% of all epochs. The number of epochs (trials) across gestures was then equalized to the gesture with the lowest number of trials.

##### Trial alignment

A custom algorithm was used to detect movement onset based on the time-frequency representation of each trial. This involved computing the short-time Fourier transform, and finding the first timepoint where the average power over channels exceeded a threshold relative to an initial baseline window (500ms). After movement onset time was identified the preceding time period was removed for each trial. The maximum allowed movement onset time was set to 1 second post gesture prompting, with the minimum being 200ms. The trial alignment was not run for Index Release and Middle Release gestures, as these immediately followed gesture offset. The final trial length was cropped to 1.5 seconds from detected movement onset for all gestures.

#### 5.3.2 Feature extraction

Feature extraction methods varied depending on the type of analysis as detailed below. In some analyses we directly used the preprocessed epoched data without further feature extraction.

##### Multivariate power frequency

For single-session classification, multivariate power frequency (MPF) features were extracted according to the method presented in (Kaifosh & Reardon, 2024). This involves computing the cross-spectral density between channels in specific frequency bins and time windows^2^, then projecting these matrices to the tangent space (Congedo et al., 2017). The following frequency bins were used: 30-62.5, 62.5-125, 125-187.5, 187.5-300 Hz. Two major deviations from the original method are: 1) computing MPF features in two equal-length (750ms) non-overlapping windows, and 2) keeping a weighted upper triangular (1 for diagonal and 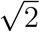 for out-of-diagonal) of the covariance matrix instead of selecting a reduced set of off-diagonals.

#### 5.3.3 Classification

We employed several classification strategies to answer differing questions. For all trainings we exclusively used OPM channels and the same 9 gestures as in (Kaifosh & Reardon, 2024): Index Pinch, Middle Pinch, Thumb Tap, Swipe Up, Swipe Down, Swipe Left, Swipe Right, Index Release, and Middle Release. We report performance for wrist only, forearm only, and for both wrist and forearm sensors.

##### Single-session

(Session-dependent) models used time-aligned preprocessed data with detected bad channels removed and MPF features were computed on this reduced channel set. The MPF features across all channels and the two non-overlapping 750 ms time windows were concatenated into a single, large feature vector on which we trained a Logistic Regression classifier. 10-fold cross-validation was employed and we report the average accuracy across the validation folds.

##### Cross-session and cross-participant

models were trained on the full channel-set to be able to match the number of features across sessions. We used the same CNN+LSTM model as in (Kaifosh & Reardon, 2024). The model was trained and tested on preprocessed data downsampled to 1000Hz without any further feature extraction. Cross-session models were run on time-aligned data, while cross-participant models were run on non-time-aligned data. Thus, the full dimensionality of a trial was 32 channels x 1500 timesteps for cross-session models and 32 channels x 2500 timesteps for cross-participant, except when reduced channel sets were used.

For cross-session analysis when using 1 session for training we set the TimeWarping segments to 3 and the max scaling for stretching to 1.5. We set the batch size to 128, the total epochs to 5000, and the patience to 800 epochs. Learning rate was set to 2e-5 and was halved every 800 epochs. When using 2 sessions for training we set the total epochs to 3000, the patience to 500, and used an initial learning rate of 5-e5, halving every 300 epochs.

In cross-participant analysis when using 20 participant for training the maximum number of epochs was set to 600, patience epochs were set to 100, and we used an initial learning rate of 1e-3, halving every 100 epochs, with a batch size of 512. When using other numbers of participants for training, we scaled the number of epochs, patience epochs, and learning-rate halving epochs with respect to the number of participants. E.g. for the 10-training-participant trainings, we set the number of epochs to 1200, and the patience and learning rate halving epochs to 200. This increase was needed due to having less training samples in each epoch.

##### Data augmentation

While time-alignment generally improves performance in the single-session case, we opted for a more robust method through data augmentation. To improve the generalizability of our model in light of data scarcity, we employed the following data augmentations. *TimeWarping* was used to deal with different gesture onset times. We randomly segmented the trial into 4 segments and applied random interpolation with a factor between 0.5 and 2, effectively shrinking or stretching the timeseries. We also employed *NoiseAugment* by adding random Gaussian noise with a 0.1 standard deviation and random rescaling of each channel with a factor between 0.5 and 2. These data augmentations were applied randomly to the training data in each batch.

##### CNN+LSTM

Similar to Kaifosh & Reardon 2024, our CNN+LSTM model had a single 1D convolutional layer with a kernel size of 20 and a stride of 5. This downsampled the input to 200Hz, which was followed by layer normalization, 3 LSTM layers, and a dense classification layer. The dense layer was applied to the last timestep output of the LSTM. The output channel number of the convolutional layer and the hidden size of the LSTM layers was set to 512. Our implementation included dropout on the channel dimension both before and after the convolutional layer with a rate of 0.2. This made the model more robust to sensor positioning differences between sessions and participants.

We trained the model with the *AdamW* optimizer with a learning rate scheduler that halved the learning rate every N epochs. We used early stopping on the validation set by ending training after M epochs (called the patience factor) have passed without improvement in validation accuracy. Batch size, learning rate, number of epochs, learning rate halving, and the patience factor varied depending on the training at hand (e.g. cross-participant vs. cross-session), due to differences in training data amounts. Thus, we report these in the respective sections. These were chosen based on manual tuning while visualizing training set performance, to achieve a smooth and relatively rapid increase in accuracy (without major fluctuations), and close to 100% final training accuracy.

All results are reported on an independent test set using the model checkpoint at the best validation accuracy in the case of deep learning trainings. We report specific data-splitting setups in the respective section in the Results. All classification was performed with custom Python code using the PyTorch package (Paszke et al., 2019).

### 5.4 Statistics

For all median accuracy values we report the inter-quantile range (IQR), which contains the middle 50th percentile of the samples in parantheses, i.e. [Q1, Q3].

For pairwise comparisons we employed the Wilcoxon signed-rank test with Bonferroni correction for multiple comparisons.

In all figures asterisks have the following meaning. one asterisk (*) denotes 1.00e-02 < p <= 5.00e-02, two asteriks (**) denotes 1.00e-03 < p <= 1.00e-02, three asterisks (***) denotes 1.00e-04 < p <= 1.00e-03, and four asterisks (****) denotes p <= 1.00e-04.

## Supplementary Results

### S.1 Single-participant classification

#### S.1.1 Gesture confusion matrices

To determine whether certain gestures were better predicted depending on the subset of selected channels, we show confusion matrices averaged across all sessions in Figure S1. We did not find any consistent and significant differences between channel subsets. The most confused gestures were Index Release and Middle Release, as they are very similar to each other and are connected anatomically by the flexor digitorum superficialis (FDS) tendons. This suggests that a hierarchical classifier which is first trained to distinguish normal gestures from release gestures and then to classify between the individual gestures could potentially have higher performance.

To verify that the two gestures were confused due to their similarity, we trained a binary classifier on the two release gestures but found the same mean accuracy as in the multiclass case (86%). However, it seems there is high variability between sessions, with session-level accuracy ranging between 54% - 99%. It would be insightful to dig deeper into why certain sessions had such poor release gesture performance. To illustrate failure modes and differences between participants, we additionally show the confusion matrices of the two lowest participants in Figure S1.

To determine whether the time window affected high and low performing participant results differently, we plotted one participant with high overall accuracy (08), and two participants with low accuracies (20 and 22) in Figure S2. It is clear that for the participant with high accuracy, the performance is already quite high across most gestures even with just a 200ms window, while participant 20 has random performance in all swipe gestures until the window size is over 500ms. In contrast, participant 22 had much worse performance in the pinch gestures at 200ms, and better performance in some swipe gestures. These results could point to differences in the timing of how these participants performed the respective gestures and could also reflect that our time-alignment algorithm may not work perfectly across all sessions. It is likely that with an improved movement onset detection algorithm the low-latency results can be much improved.

### S.2 Cross-session classification

Confusion matrices for cross-session trainings with 2 training sessions are shown in Figure S3. Release gestures seem to be the most confused, followed by swipe right with swipe up. In addition to aggregating over all participants we show the confusion matrices aggregated over the 3 participants with highest and lowest cross-session accuracy, respectively. Gestures that are particularly confused for low performance participants include the index pinch, thumb tap, swipe up, and the release gestures. Using forearm-only channels better performance is observed in the two swipe gestures. We also observed instances in which a single session had poor cross-session performance (bottom row of Figure S3), suggesting that specific sessions may have very different characteristics compared to other sessions.

### S.3 Cross-participant classification

The confusion matrices across all test participants are shown in Figure S4. Gestures that are particularly confused are the swipe up,swipe right, and swipe down. Compared to single-session and cross-session trainings the release gestures have much higher accuracy in cross-participant trainings. We also show confusion matrices of 14 sessions with highest and lowest test accuracy, respectively. Low-accuracy sessions seem to have higher confusion rates in the two pinch gestures and the swipe down.

### S.4 Latent space visualization

To visualize the degree of variability present in the data we trained Uniform Manifold Approximation and Projections (UMAPs) on the datasets using the MPF features of each trial (Supplementary Figure S5). We found visible clustering when trained with a single dataset which degraded as we used additional datasets (Supplementary Figure S5a), demonstrating the decrease in classification accuracy when introduced with generalization.

To quantify the differences we calculated the silhouette score, or how well the gestures cluster in latent space, of a UMAP trained on a single session then the silhouette score of all other datasets projected on that UMAP (Supplementary Figure S5b). There were significant differences for each gesture, even Index and Middle releases that were typically poorly classified.

### S.5 Single-session and cross-participant model accuracy correlations

To assess whether the size of the participants’ wrist and forearm, number of bad trials, or number of bad channels affected single-session or cross-participant accuracy we show them in Supplementary Figure S6. There was no significant correlation with wrist and forearm size and model accuracies. The number of bad trials and bad channels both were significantly correlated with single-session accuracy (p<0.05, Spearman’s rank correlation coefficient), but they were not correlated with the cross-participant accuracy. Finally, single-session and cross-participant accuracies were not correlated with one another.

### S.6 Pairwise channel coherence

We calculated pairwise coherence between adjacent channels, channels across the band, channels located on the same position between the two bands, as well as channels in the same sensor (i.e. tangential and radial components at the same location) to determine the similarity between channels (Supplementary Figure S7. Coherence between two channels, *C*_*xy*_, was calculated using magnitude squared coherence which effectively shows similarity at different frequencies:

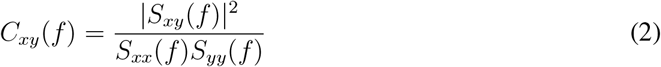

where *S*_*xy*_ is the cross spectral density between the two signals, and *S*_*xx*_ and *S*_*yy*_ are auto spectral densities of each signal. Channels adjacent from one another had the highest coherence values mostly throughout the bandwidth. Channels across the bands had the highest coherence at lower frequencies, likely as they are directed in the same axis. The coherence decreased with higher frequency, showing that higher frequencies reflect local activity. The wrist and forearms had very different information, as they are likely detecting different muscle groups. Finally, the two axes of the same sensor had lower low frequency coherence but high coherence overall, likely due to crosstalk inherent between channels in OPM sensors.

### S.7 Variability in task performance

To both track participant performance as well as consistency of each trial we extracted joint positions from simultaneous recordings with the Leap Motion Controller. Supplementary Figure S8 shows joint positions of all Swipe Right trials for a single session for two different participants. We found large variability in both within session and between participant trial performance in finger positions as well as hand angle and orientation. Though expected, this suggests user training will also help improve classification accuracy.

**Supplementary Figure S1:**
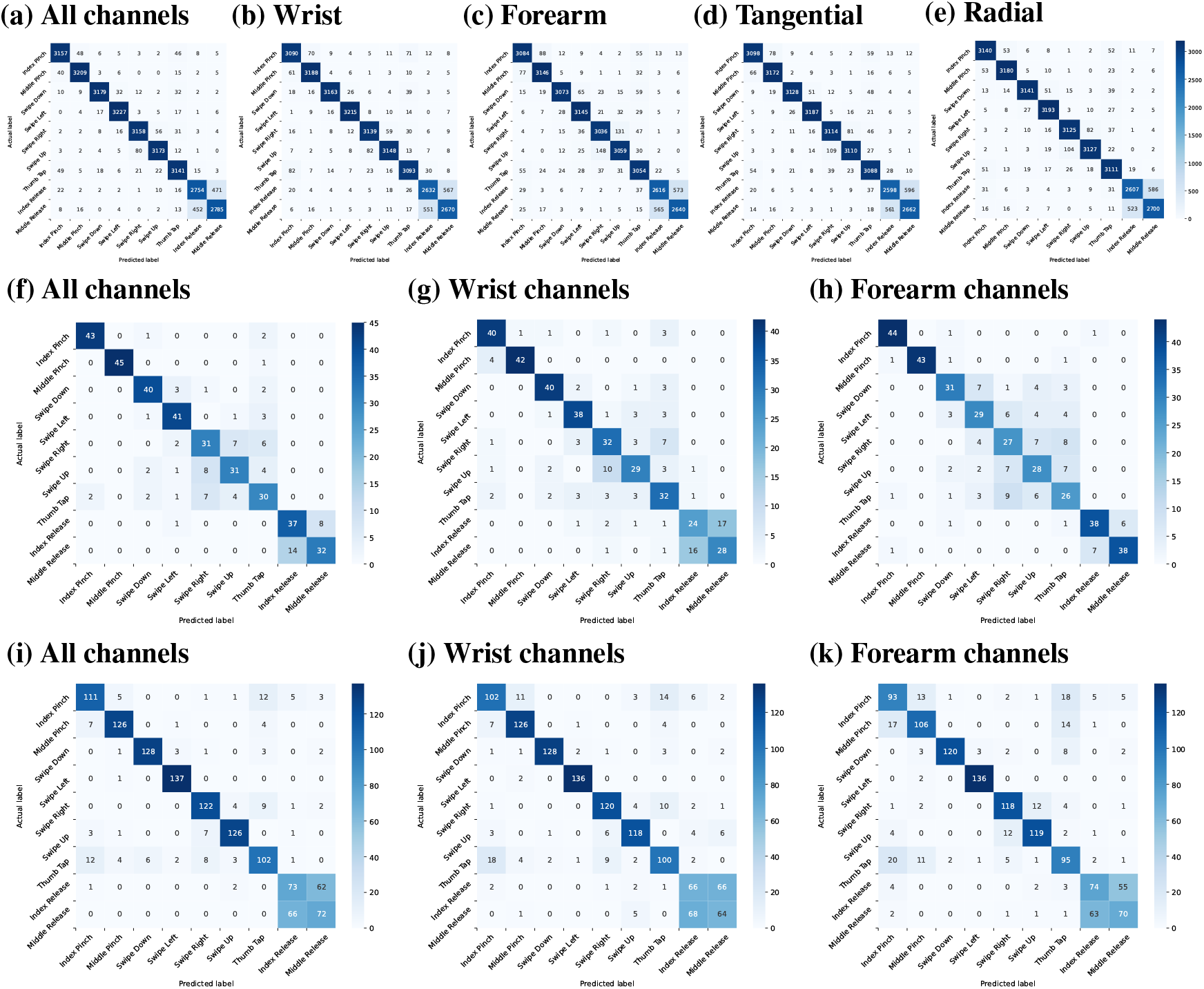
Investigation of single-participant confusion matrices. (a-e) Confusion matrices across sessions for various channel subsets. Index and Middle Release are the most confused gestures, and there are no interesting differences between the channel subsets. (f-k) Confusion matrices of 2 participants with low performance (rows). (f-h) - participant 20, (i-k) - participant 22. We sought to better understand why certain sessions and participants had low accuracy. To this end, we show the confusion matrices across all sessions of participants 20 and 22, which both had mean accuracies of 80%, and 80% using all channels. It is interesting to note that participant 22 has close to 50% accuracy in the two release gestures, while in participant 20, the swipe gestures are confused more. There are also differences depending on whether wrist or forearm channels are used in classification. The release gestures are better classified for participant 20 with forearm-only channels, but swipe gesture accuracy is lower.

**Supplementary Figure S2:**
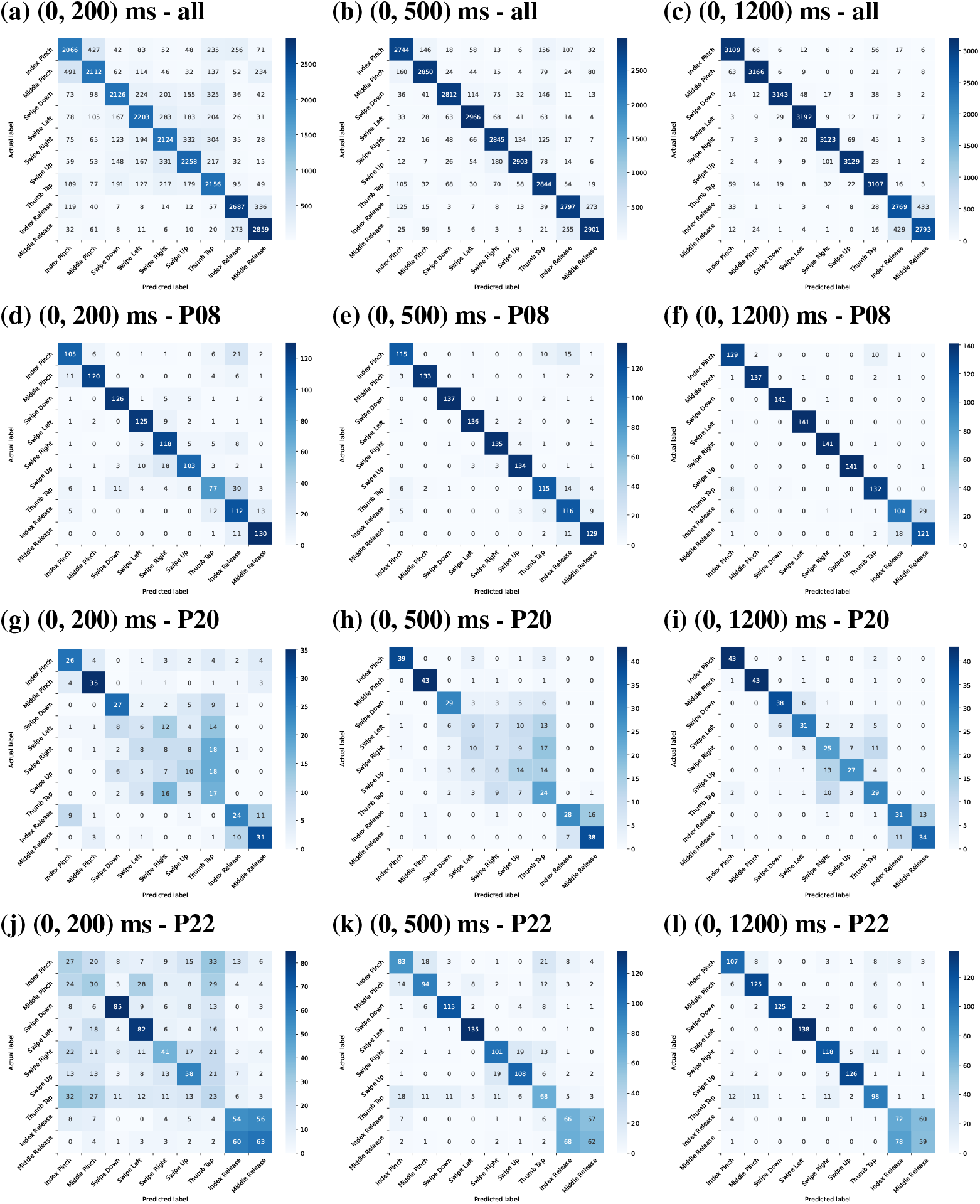
Confusion matrices for 3 time windows. (a-c) Confusion matrices across all sessions for 3 time windows. Interestingly, release gestures are better classified using a smaller window compared to the full trial length. Pinch gestures seem to be most affected by a reduced window length. (d-l) Confusion matrices of individual participants (rows). (d-f) - participant 08 (high accuracy), (g-i) - participant 20 (low accuracy), (j-l) - participant 22 (low accuracy). While participant 08 already has high accuracy with a 200ms window, this is not the case for the two participants with low accuracies, and the confusion matrices show that there are differences between them in terms of which gestures are poorly classified with a small window.

**Supplementary Figure S3:**
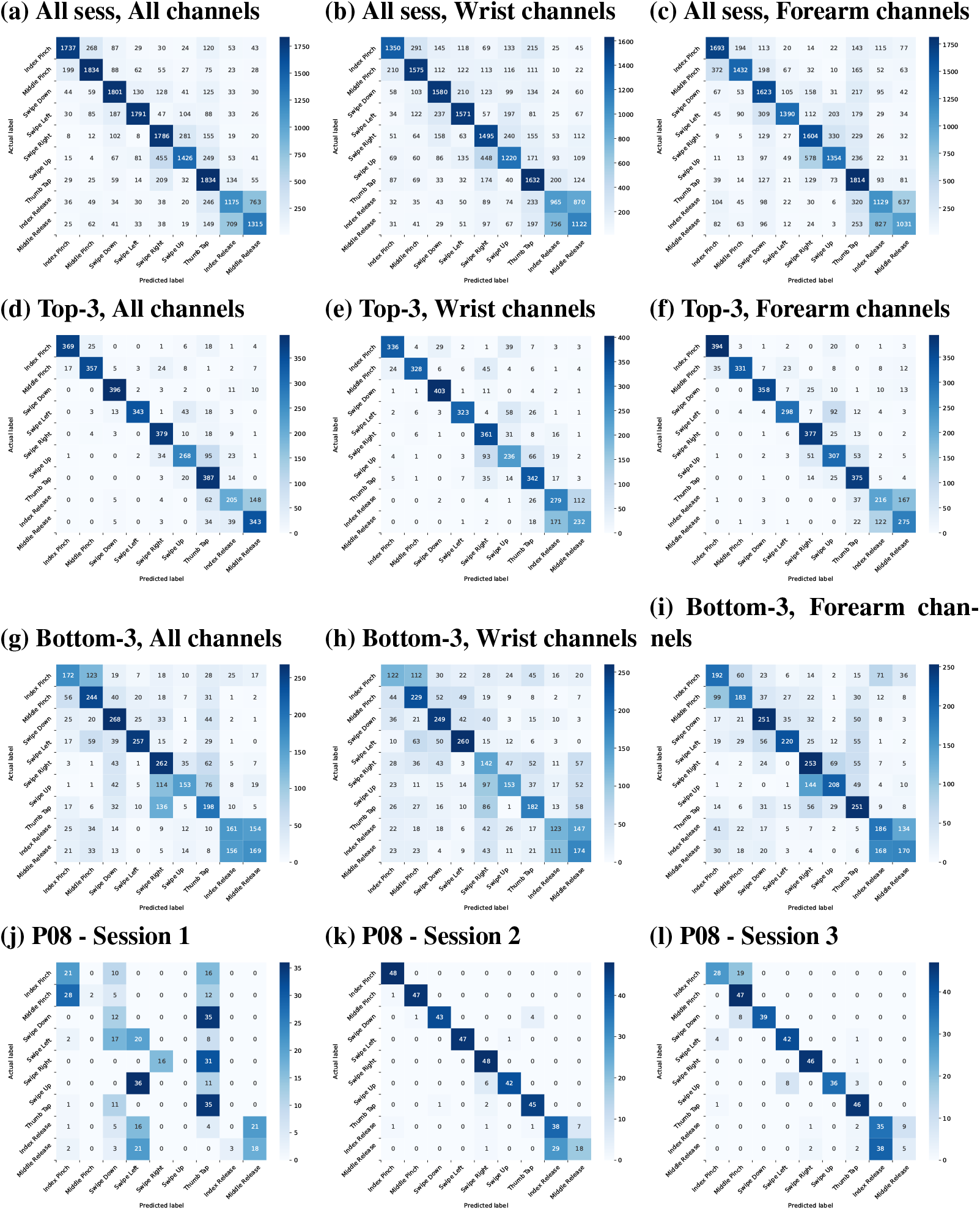
Cross-session confusion matrix investigation. (a-i) Confusion matrices for all participants (a-c) the 3 highest accuracy participants (d-f) and 3 lowest accuracy participants (g-i). Release gestures are most confused, followed by swipe up gesture. 2 training sessions were used in each case, and the confusion matrices are aggregated over all test sessions. The top-3 and bottom-3 participants were selected based on the all-channel performance. (i-l) Confusion matrices for the 3 (test) sessions of participant 08. Session 2 has much lower performance when being tested on compared to the other sessions. 2 training sessions were used in each case.

**Supplementary Figure S4:**
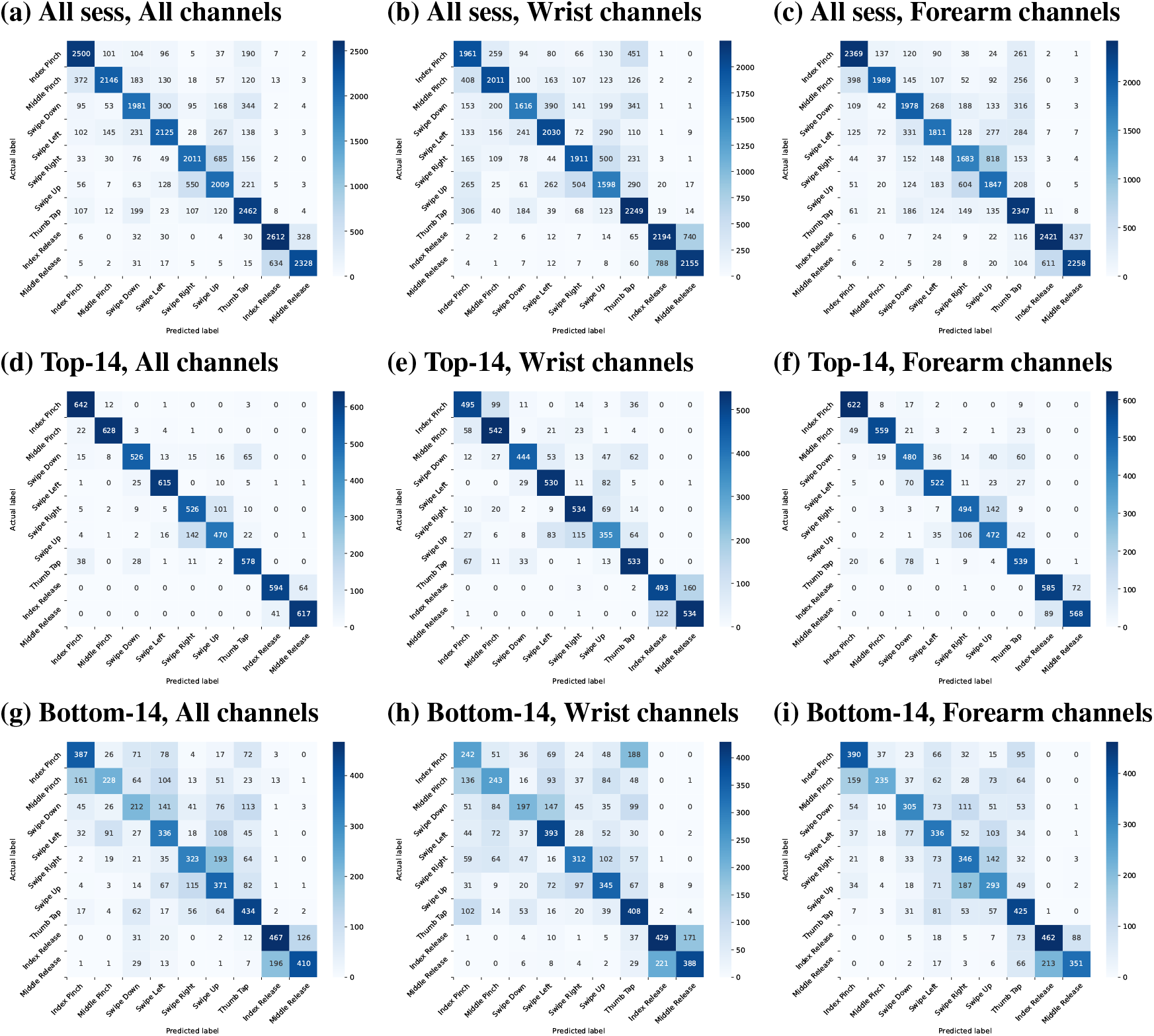
Cross-participant confusion matrices for all sessions. (a-c) the 14 highest accuracy sessions (d-f) and 3 lowest accuracy sessions (g-i). Low accuracy sessions have higher confusion rates in the two pinch gestures and the swipe down. The top-14 and bottom-14 sessions were selected based on the all-channel performance.

**Supplementary Figure S5:**
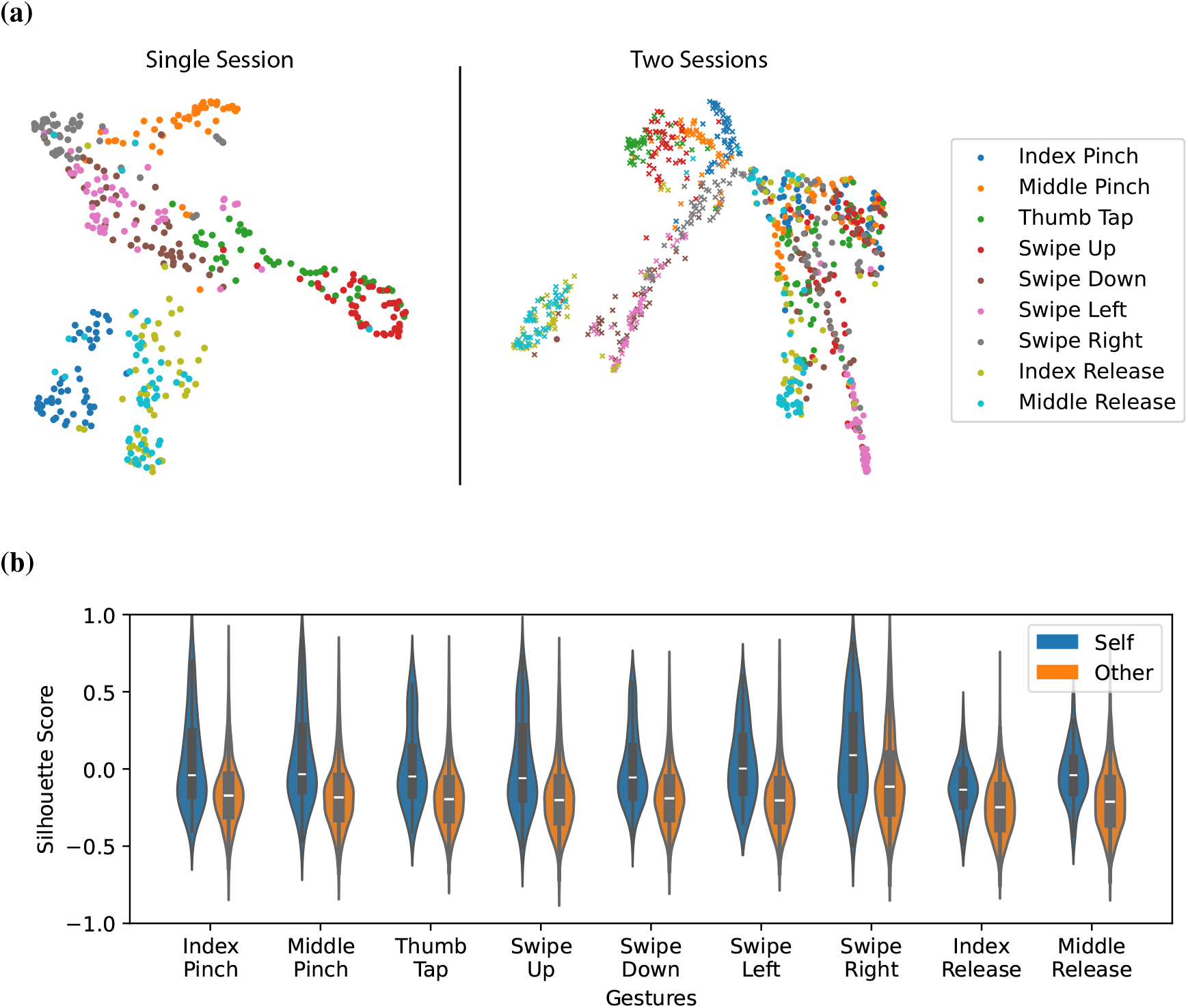
Latent space analysis. (a) Latent space calculated using Uniform Manifold Approximation and Projection (UMAP) of a single session (left) and two sessions combined (right) – the different marker shapes denote the two separate sessions. MPF features of the full 2.5 trial duration was used. Notice the clear clustering between the different gesture types in the single session UMAP. The clustering is degraded as additional sessions are included, similar to difficulties in generalizing classification. (b) Silhouette scores showing how well the gestures are clustered when training a UMAP with the dataset (Self) compared to projecting the data onto a UMAP trained on another dataset. The silhouette score spans from -1 (worst clustering) to 1 (best clustering). Projections onto other UMAPs provides worse clustering for each gesture type demonstrating variability between sessions and subjects. Notice the Index and Middle Release gestures are not well clustered even in the Self condition.

**Supplementary Figure S6:**
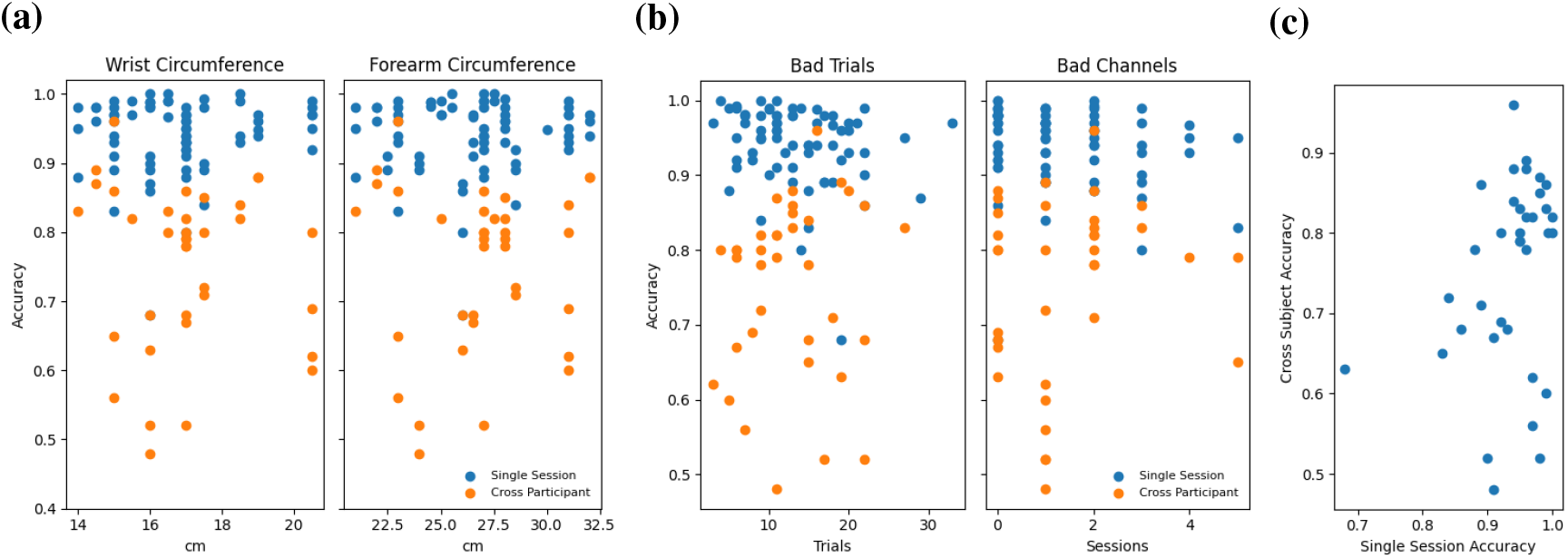
Single- and cross-participant accuracy vs various metrics. Comparisons of single-session model accuracy and cross-participant model (using 20 training participants) accuracy with a) wrist and forearm circumference of the participant, b) number of bad trials and bad channels in each session, and c) between one another. Single-session accuracy is statistically correlated with the number of bad trials and number of bad channels (p<0.05) but has no significant relationship with any other metric. Cross-participant accuracy is not correlated with any metric. Single-session and cross-participant accuracies are not correlated with one another.

**Supplementary Figure S7:**
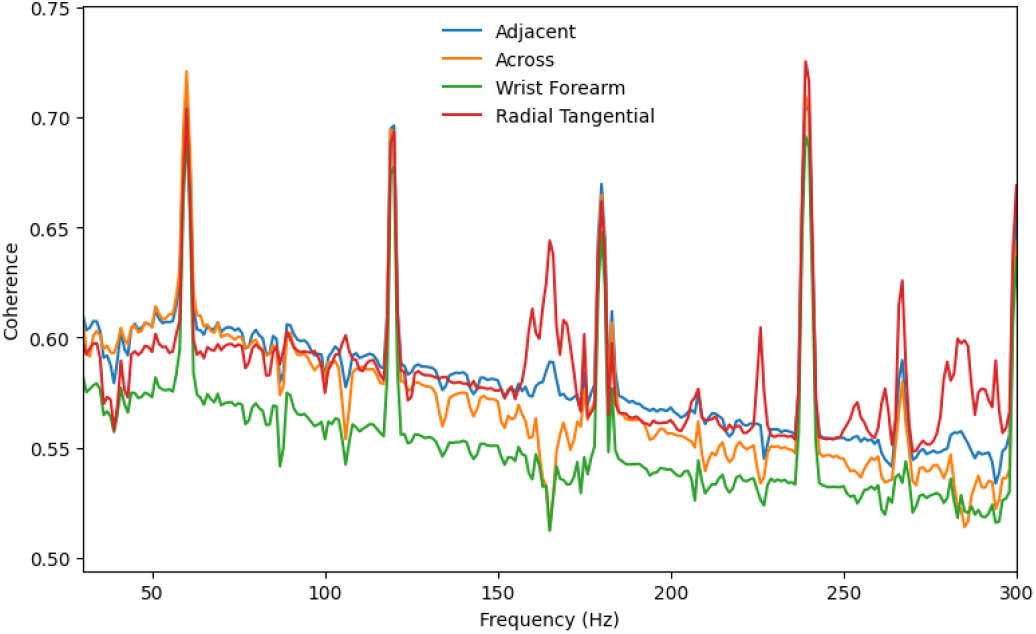
Pairwise channel coherence. Coherence calculated between channels adjacent from one another (blue), across the band from one another (orange), one the same position between the wrist and forearm (green) and within the same sensor (red). For the first three, all pairs were of the same axis for each channel (i.e. radial with radial, tangential with tangential).

**Supplementary Figure S8:**
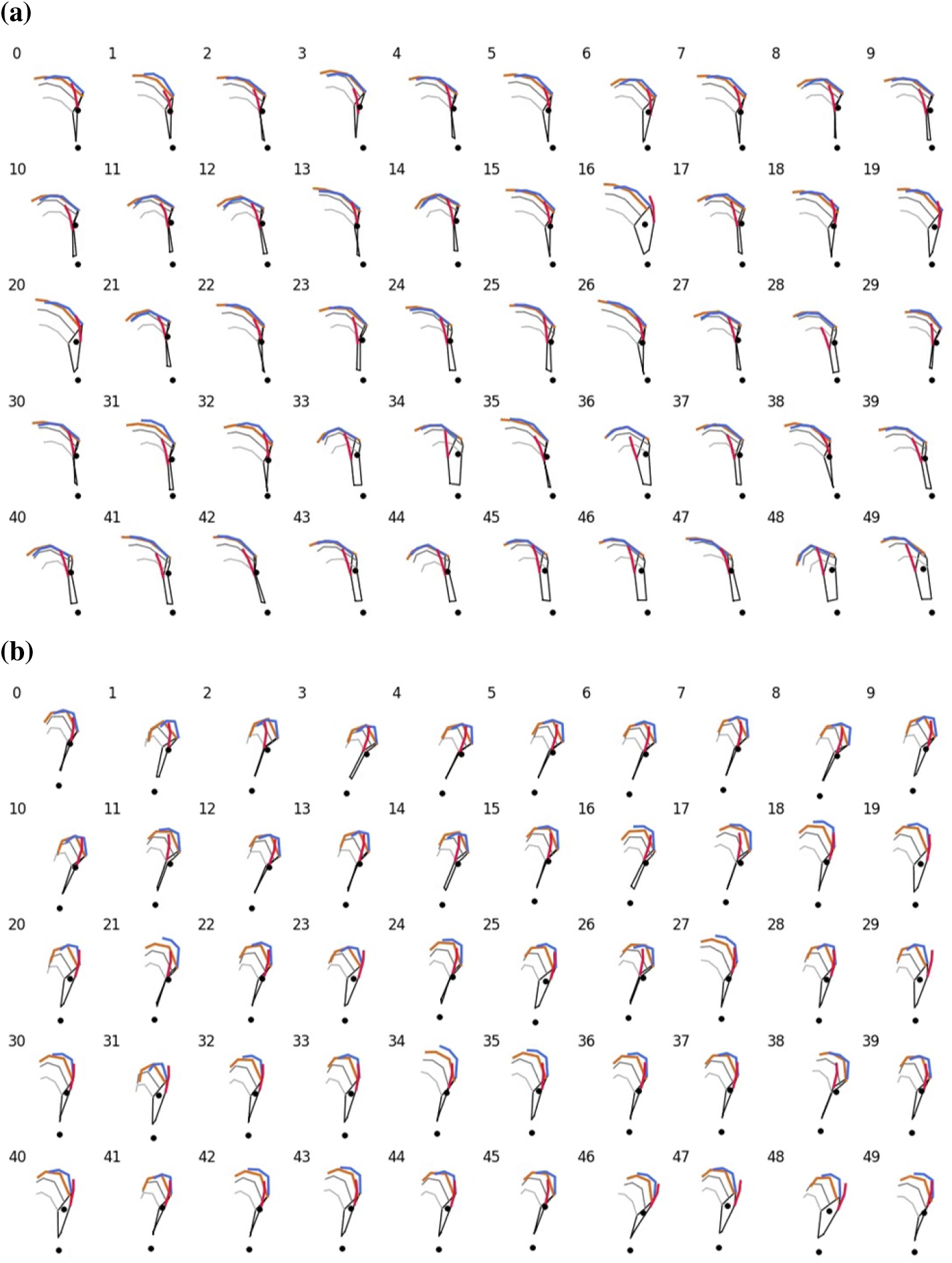
Variability of task performance. Hand and finger joints tracked with the Leap Motion Controller during all Swipe Right trials in two sessions of two different participants 2.5 seconds after cue presentation (a, b). Notice the variability in hand and finger positions within session and between participants.

## Footnotes

1 https://github.com/OHBA-analysis/osl

2 https://github.com/pyRiemann/pyRiemann/tree/master?tab=readme-ov-file

